# Mapping Whole-Brain Auditory Activation with 3T Multi-Echo fMRI at the Group and Individual-Subject Level

**DOI:** 10.64898/2025.12.03.692160

**Authors:** Michelle C. Medina, Neha A. Reddy, Kevin R. Sitek, Molly G. Bright

## Abstract

Magnetic resonance imaging (MRI) is a powerful and established tool to non-invasively probe the human auditory system. Varied blood oxygen level-dependent functional MRI (BOLD fMRI) acquisitions have been used to examine the functional roles of this system, but these acquisitions have substantial limitations, such as the need for specialized hardware and long acquisition times, and they typically rely on group averaging of activation patterns. In recent years, whole-brain multi-echo (ME) fMRI techniques have been used to reduce artifacts and scan times, map entire sensory systems, and improve sensitivity to neural activity in both resting-state and task fMRI data acquired at 3T. Combined with dense-sampling strategies, these ME techniques have facilitated “precision mapping” of neural activity in individual subjects. Thus, in this technical note we propose the use of a commonly available ME wholebrain acquisition and ME denoising approaches to examine the auditory system in both group and densely sampled single-subject datasets. Whole-brain and region-specific analyses were performed to identify auditory regions of activation. At the group level, auditory activation was identified bilaterally in cortical regions and unilaterally in cerebellar lobules VIIb/VIIIa with both analyses. Additionally, the region-specific analysis successfully identified unilateral activation in thalamic and brainstem regions. At the individual subject-level, precision mapping combined with ME denoising methods enhanced sensitivity, yielding bilateral activation in cortical, cerebellar, thalamic, and brainstem regions with both analyses. Lastly, we demonstrate the benefits of using multi-echo methods and a whole-brain precision mapping approach to better align an individual’s functional response to their specific anatomy.

## 1. Introduction

Magnetic resonance imaging (MRI) is a powerful and established tool to non-invasively probe the human auditory system. In recent years, the emergence of high-field 3T and ultra-high-field 7T scanners has produced exquisite anatomical delineations of this system’s prominent structures, yielding detailed atlases of both cortical and subcortical auditory regions^1,2^. Furthermore, the use of blood oxygen level-dependent functional MRI (BOLD fMRI) has allowed the investigation of these structures’ functional roles (e.g. sound detection^3^ and auditory selective attention^4^) and their responses to various sound stimuli^5–9^.

Although varied fMRI acquisitions have been used to examine auditory function^10–13^, these acquisitions have substantial limitations. Many rely on specialized hardware and elaborate sequences with limited availability, which may also require long acquisition times. To address signal-to-noise (SNR) challenges and the small size of subcortical auditory nuclei, fMRI of these regions is typically performed with ultra-high-field scanners, which can improve sensitivity and spatial resolution but are yet to be widely accessible. Alternatively, a high-resolution reduced field-of-view acquisition can be employed with more conventional 3T scanners, focusing on specific thalamic or brainstem nuclei rather than capturing the entire auditory system at once^3,6^. SNR is also frequently improved by averaging together functional maps across individuals in a group-level statistical analysis^10^, where misalignment of small nuclei during image coregistration^1^ can impact sensitivity. These challenges make it difficult to reliably and holistically investigate human auditory pathways and processing.

Whole-brain multi-echo fMRI techniques have been used to reduce artifacts and improve sensitivity to neural activity in resting-state and task fMRI data acquired at 3T. Recently, such techniques have been applied to successfully map the supraspinal tactile sensory system, identifying activation in cortical, subcortical, cerebellar, and brainstem regions in one wholebrain acquisition^14^. Furthermore, multi-echo fMRI has been applied in “precision mapping” experiments^15–17^, in which dense-sampling of individuals can generate activation maps in single subjects, which may avoid the challenges of co-registering small subcortical nuclei across individuals and provide individual-specific functional localization. In this technical note, we applied this systems-level, multi-echo, dense-sampling fMRI approach to the auditory system, demonstrating a robust strategy for characterizing activity at both the group and individual-subject level.

## 2. Rationale

In multi-echo (ME) fMRI, two or more image volumes are collected for each time point during data acquisition, each reflecting a different sensitivity to BOLD contrast, and subsequently combined using a weighted average. This “optimal combination” dataset benefits from the higher signal in the volumes collected at earlier echo times, boosting SNR, while remaining highly sensitive to BOLD contrast (and thus neural activation) from later echo times^18^. Additionally, the signal decay across echo times can be examined in combination with techniques such as independent component analysis (ME-ICA) to differentiate BOLD-related signals from non-BOLD artifacts (e.g., head motion), facilitating data denoising.

The implementation of ME-ICA has become increasingly common in recent years given its readily available protocols, easy-to-use software, and growing translational impact^17,19^. Previous studies have shown its benefits in mitigating task-correlated motion in healthy and patient populations^20,21^ as well as in enhancing task-based BOLD sensitivity and specificity across entire neural systems at clinically available field strengths^14,22^. Moreover, ME-ICA has facilitated the success of precision mapping studies, reducing the total scan time needed to yield reliable results^16,17^.

In summary, ME methods can be used to successfully acquire whole-brain fMRI data in standard 3T MRI scanners, improve data quality by boosting sensitivity in challenging regions, and robustly capture both group-level and single-subject information. Thus, in this technical note we propose the use of a commonly available ME whole-brain acquisition and ME-ICA approaches to examine the auditory system in both group and densely sampled single-subject datasets.

## 3. Methods

### 3.1 Data Collection

Fourteen healthy individuals (23±1y, 5F) with no known history of neurological, vascular, or auditory disorders were enrolled in the study after providing written, informed consent. The study protocol was approved by the Institutional Review Board of Northwestern University. Participants were scanned using a Siemens 3T Prisma and 32-channel head coil. All fourteen individuals participated in Session 1, which included an anatomical scan and one functional MRI run. A multi-echo T1-weighted MPRAGE was collected and used for registration^23^: TR = 2.17 s, TEs = 1.69/3.55/5.41 ms, TI = 1.16 s, FA = 7°, voxel size = 1 x 1 x 1 mm^3^, FOV = 256 x 256 m^2^. The three echo MPRAGE images were then combined using root-mean-square. A functional scan was collected using a multi-band, multi-echo gradient-echo echo-planar imaging sequence with whole-brain coverage provided by the Center for Magnetic Resonance Research (CMRR, Minnesota)^25,26^: TR=2.2s, TEs=13.40/39.5/65.6ms, MB factor=2, voxel size=1.731×1.731×4mm^3^, 44 slices, FOV=180mm, and 250 volumes^14,25,26^. Axial slices were aligned perpendicular to the base of the fourth ventricle to align with brainstem anatomy while achieving maximum whole-brain coverage. A reverse phase encode functional image was collected and used for distortion correction^24^. Three participants (23±1y, 2F) returned for a “precision mapping” session (Session 2) to collect four additional functional scans. During all scanning, end-tidal CO_2_ was measured using a nasal cannula and gas analyzer system at a sampling rate of 1000 Hz (PowerLab, ADInstruments).

### 3.2 Auditory Setup and Stimulus

Avotec’s Silent Scan system and Sensimetric S14 earbuds with disposable foam canal ear tips (Comply) were used to transmit the audio with an average noise reduction rating above 29 dB. MULTIPAD EAR inflatable pads (Pearl Technology) were placed between the head coil and the patient for additional comfort and immobilization. Participants were instructed to attentively listen to the music, remain as still as possible, and stay awake while looking at a fixation cross. During each functional scan, participants listened to the auditory stimuli in a block design: 10-seconds of an RMS-normalized, instrumental-only pop song was followed by 15-seconds rest for a total of 21 trials, with the exception of two participants who had 20 trials. Three different pop songs were used to maintain attention, and song choice was randomized across participants and scans.

### 3.3 Data Analysis

#### 3.3.1 MRI Pre-processing

All pre-processing steps were performed using FSL^27^ (version 6.0.3) and AFNI^28^ (version 24.1.11). T1-weighted images for all participants were processed, including bias field correction and brain extraction (*fsl_anat*). The first 10 volumes of each functional scan were removed for the signal to reach steady-state equilibrium. Motion realignment parameters were calculated using the first echo data and the Single Band reference image (*3dvolreg*) and applied to all echoes (*3dAllineate*). Brain extraction (*bet*) and distortion correction (*topup, applytopup*) were performed on all images.

#### 3.3.2 Multi-echo with independent component analysis (ME-ICA)

Echo timeseries were optimally combined^29^ and ICA decomposition was performed using tedana^19^ (version 24.0.2). Rejected (i.e., noise-related) ICA components were automatically classified using tedana’s external regressors decision tree and orthogonalized using the conservative approach in Moia et al^21^. ME-ICA data were smoothed using a 3 mm FWHM Gaussian blur (*3dMerge*) and converted to signal percent change.

#### 3.3.3 General Linear Model

The auditory stimulus’s timing was convolved with the canonical double-gamma hemodynamic response function to generate the auditory task regressor. Subject-level and precision mapping concatenated results were modeled using a general linear model (GLM) and processed using AFNI’s *3dREMLfit*. The ME-ICA model included six motion regressors and their derivatives, HRF-convolved end-tidal CO_2_, the auditory task regressor, the rejected ICA components and up to fourth-order polynomial terms.

#### 3.3.4 Group-level Analyses

Beta-coefficient and t-statistic maps were registered to 2-mm Montreal Neurological Institute (MNI) space (*FNIRT, FSL*). Group-level activation maps were obtained by performing a wholebrain bi-sided one-sample t-test (*3dMEMA, AFNI*), thresholded at *p* < 0.005 and clustered at *α* < 0.05 (*3dFWHMx, 3dClustSim, 3dClusterize, AFNI*). As region-specific analysis has been previously conducted to better elucidate activation in challenging subcortical regions^14,30^, we employed this methodology to boost sensitivity in regions known *a priori* to be involved with auditory processing: thalamus, cerebellum, and brainstem. The group-level region-specific analysis was performed using a one-sample t-test (*RANDOMISE*^31^) with threshold-free cluster-enhancement^32^ and family-wise error correction (*p_c_* < 0.05) within masks of each region. The thalamus and brainstem masks were created by thresholding these regions from the HarvardOxford Subcortical Structural Atlas^33^ at 50% and 25%, respectively. Masks were manually edited to ensure coverage of the medial geniculate nuclei (MGN) and inferior colliculi (IC) using the subcortical auditory atlas provided by Sitek et al^1^. The cerebellar mask was created by thresholding the MNI structural atlas^34,35^ region at 50%.

#### 3.3.5 Precision Mapping Analyses

In the three individuals who underwent repeated “dense-sampling” scanning, the four functional scans from the second session were concatenated prior to subject-level modeling. To evaluate the impact of scan length on activation results, this was done with subsets of the data, considering the first 10, 20, 30 or all 40 minutes of scanning.

A single-echo (SE) model was additionally implemented to assess if ME offered benefits in identifying auditory nuclei at the individual-subject level. The second echo (39.5 ms) data were used as a surrogate for conventional SE analysis. The SE data were smoothed using a 3 mm full width at half maximum (FWHM) Gaussian blur (*3dMerge*) and converted to signal percent change. The SE model included all the regressors included in the ME-ICA model with the exception of the rejected ICA components.

Significant voxels of positive activation were found in a whole-brain mask in functional space, with False Discovery Rate correction (*p_c_* < 0.05). To better localize the areas of most dominant activation, a region-specific analysis was performed by extracting voxels with the top 5% t-statistics within the same thalamus, cerebellum, and brainstem masks detailed in the previous section transformed to each subject’s functional space. These analyses were conducted for each densely sampled subject using their 10, 20, 30, and 40-minutes concatenated data for both SE and ME-ICA models.

#### 3.3.6 Comparison of Group-Level and Single-Subject Activation

A comparison between the region-specific group-level results and each of the ME-ICA precision mapping results (using all 40 minutes of scan data) was performed by overlaying the activation clusters for both approaches in standard and individual anatomical space.

#### 3.3.7 Comparison of SE and ME Precision Mapping Results

To assess spatial overlap between activation clusters from the SE and ME-ICA region-specific results, dice similarity coefficients were calculated. For each region, the top 5% t-statistics cluster from the 40-minutes concatenated ME-ICA data was used as the ground truth. Although this is not a true “ground truth” as such, these ME-ICA results fully included the unilateral activation detected in the SE data, as well as additional bilateral regions expected *a priori* to be activated by our stimulus. Thus, results from the 40-minutes concatenated ME-ICA data were compared with results from activation clusters from all remaining ME-ICA and SE data subsets. All subject-specific maps and activation results were then transformed to each subject’s anatomical space for visualization.

## 4. Results

### 4.1 Group-level Activation

Significant clusters of activation were mapped bilaterally in primary and secondary auditory cortices and unilaterally in cerebellar lobules VIIb/VIIIa in both whole-brain and region-specific analyses (***Figure 1***). Although the cerebellum has been primarily known for its involvement in motor control^36^, recent evidence suggests it might play a significant role in cognitive functions^37^, such as auditory processing^38^ and sound discrimination^39^. Indeed, various studies have consistently shown lateralized activation of the cerebellar left Crus I area during pure tones and clicks passive auditory tasks^38,39^. Furthermore, lobules VIIb/VIIIa have been associated with visual attention^40^, with the left lobule VIII having a prominent involvement during music tasks^37^.

**Figure 1:**
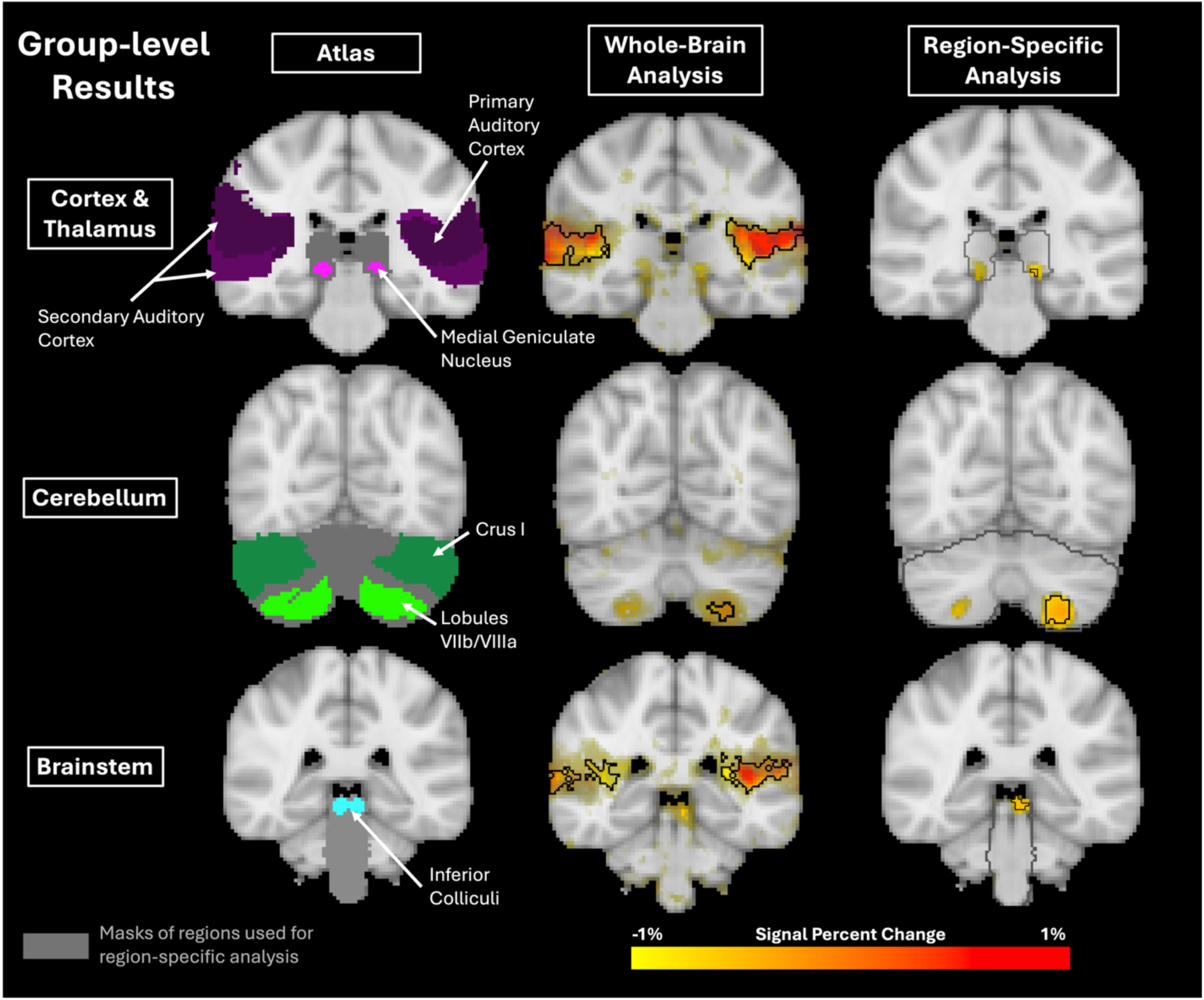
Group results for whole-brain (3dMEMA, bi-sided t-test, thresholding at *p* < 0.005, clustering at *α* < 0.05) and region-specific (RANDOMISE, one-sided t-test, TFCE and FWE *p_c_* < 0.05) analyses. The opacity of the beta-coefficient values was modulated by t-statistic. Black outlines showcase significant clusters of positive activation, compared to rest. Atlas regions are shown for comparison. Grey masks and outlines denote the regions used for the region-specific analysis.

The thalamic MGN and brainstem IC regions are well-established components of the auditory system. Activation of these regions is present in the whole-brain analysis maps, but remains subthreshold (i.e., not statistically significant). The region-specific analysis, however, shows significant unilateral activation in the left MGN and left IC. Subthreshold activation on the right hemisphere is present in MGN and cerebellar regions, but not in IC.

### 4.2 Precision Mapping Activation

For all densely sampled subjects, the whole-brain analysis reveals bilateral areas of significant positive activation in the primary and secondary auditory cortices, MGN, cerebellar lobules VIIb/VIIIa, and IC (*Figure 2*). A region-specific analysis was performed to better localize the most dominant areas of activation within each region expected to be involved with the auditory system. Note the similarity between the clusters of positive activation present in the whole-brain and region-specific analyses.

**Figure 2:**
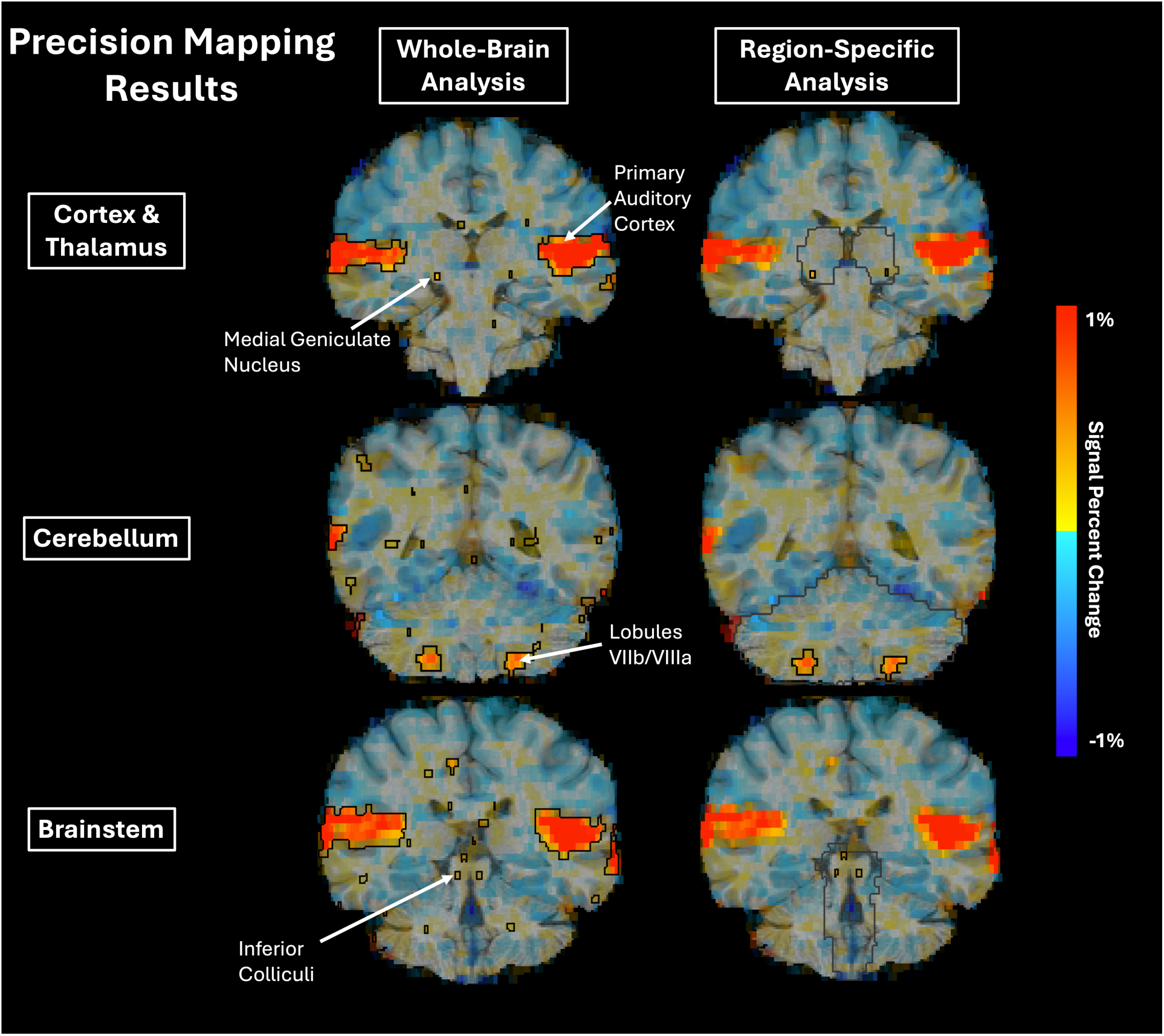
Single-subject results for whole-brain (FDR *p_c_* < 0.05) and region-specific (top 5% tstatistics) analyses for a representative subject in anatomical space. The opacity of the betacoefficient values was modulated by t-statistic. Black outlines showcase clusters of activation. Grey outlines denote the regions used for the region-specific analysis.

### 4.3 Comparison of Group-Level and Single-Subject Activation

To better understand the differences between the group-level and precision mapping regionspecific results, the activation clusters for both approaches were overlayed in standard and individual anatomical space (*Figure 3*). In standard space, subject-specific clusters show variable locations within regions. Minimal overlap is present in the MGN area, and modest overlap is seen in lobules VIIb/VIIIa and IC regions. This result is perhaps unsurprising given that the thalamus is known to suffer from poor MR contrast, hindering the differentiation between its substructures^41–43^ and further exacerbating registration challenges in this region^43–45^. Although various atlases^43,46^ and thalamus-specific registration algorithms^45,47^ have been proposed to improve the identification and alignment of small thalamic nuclei, these tools are yet to be tailored for the additional registration difficulties that may arise with a whole-brain functional acquisition, such as the one presently used. Additionally, previous studies report low anatomical overlap of auditory subcortical structures between subjects^1^, even at higher field strengths with better spatial resolution, suggesting that inter-individual variabilities are also a likely source behind these discrepancies.

**Figure 3:**
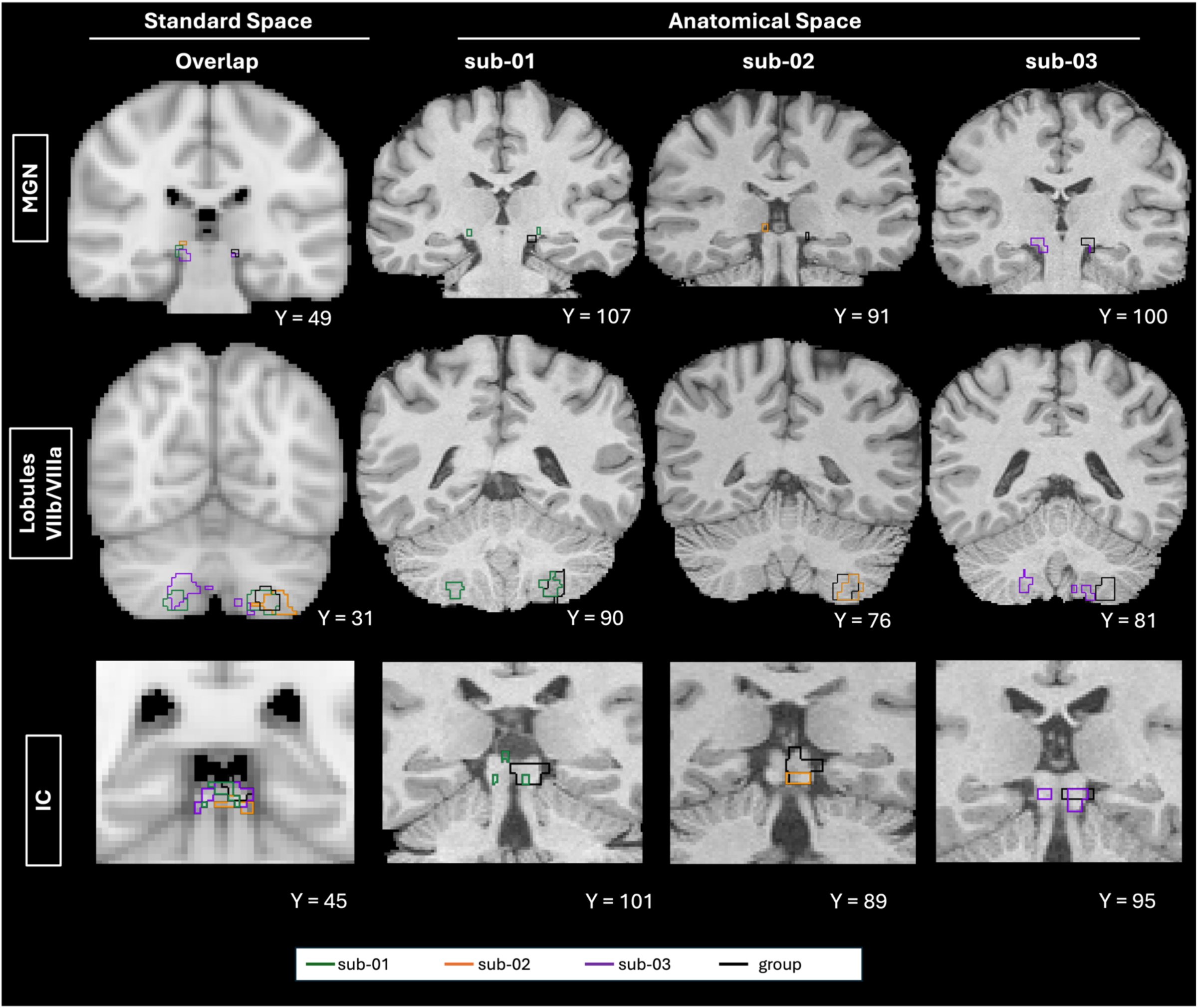
Overlayed group-level and precision mapping region-specific results in standard and anatomical space. Group-level activation clusters are shown as black outlines and precision mapping activation results are depicted as colored outlines.

Despite these challenges, the precision mapping approach offers improved alignment between an individual’s functional activation and their own anatomy, as seen when projecting on each subject’s anatomical space. Activation clusters are located within the confines of the expected anatomical structures across all subjects and regions, showcasing the advantages in specificity gained by using a precision mapping approach, which could be especially critical in clinical cohorts when trying to understand an individual’s unique impairment.

### 4.4 Comparison of SE and ME Precision Mapping Results

Lastly, to investigate if ME techniques yielded any benefits in identifying auditory nuclei, we compared conventional SE and ME-ICA models across all densely sampled subjects by conducting a spatial overlap assessment between their region-specific activation results. Dice coefficient scores for the ME-ICA model increased with increasing scan duration in all but one subject (sub-02) across all regions (*Figure 4A*). (NB: a similar plot for the cerebellum can be found in ***Supplemental Figure 1***). The dice coefficient scores for the SE model presented variable trends with increasing scan duration across subjects and regions. For every subject and across all regions, ME-ICA yielded equivalent or higher Dice coefficient scores than SE starting at 20 minutes of data. In regions such as the thalamus and brainstem, with small auditory nuclei with poor contrast and physiological noise, ME-ICA succeeded in mapping bilateral activation clusters in both MGN and IC regions starting at 30 minutes of data (*Figure 4B*). In contrast, SE maps required all 40 minutes of data to map the bilateral MGN cluster while the IC cluster remained unilateral even with all 40 minutes of data, as shown by the dashed pink and blue arrows.

**Figure 4:**
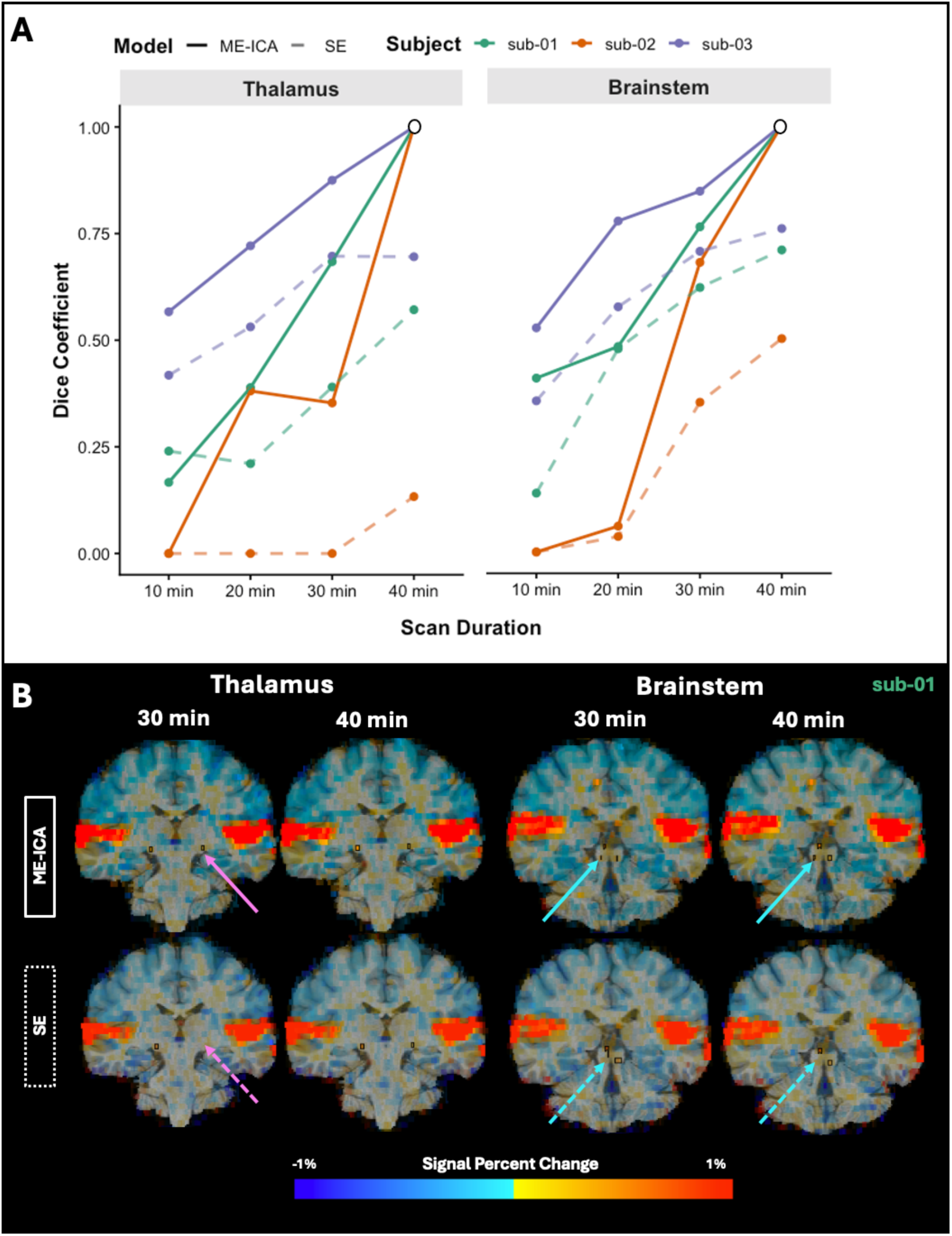
Comparison between ME-ICA and SE models. (A) Dice similarity coefficient plots between ME-ICA (solid line) and SE (dashed line) region-specific activation results across scan duration. The 40-minute concatenated ME-ICA top 5% t-statistics cluster was used as groundtruth; thus, a blank circle was overlayed with each subject’s 40-minute scan duration Dice coefficient value. Each colored line represents a precision mapping subject. (B) Precision mapping results for region-specific analysis using ME-ICA and SE models for a representative subject. The pink and blue solid arrows indicate the bilateral activation identified by ME-ICA for MGN and IC regions, respectively. The pink and blue dashed arrows show the missing left and right activation cluster for MGN and IC regions, respectively, using the SE model.

## 5. Conclusions

We present a whole-brain fMRI investigation of the auditory system using a 3T field strength scanner and novel multi-echo methods. Whole-brain and region-specific analyses were performed in both group-level and precision mapping data to identify auditory regions of activation. At the group level, auditory activation was identified bilaterally in cortical regions and unilaterally in cerebellar lobules VIIb/VIIIa with both analyses. Despite common challenges such as imperfect anatomical co-registration, inter-individual differences in functional anatomy, and low SNR and poor contrast in the auditory regions of the brainstem and thalamus, a regionspecific analysis successfully identified unilateral activation in the MGN and IC. Furthermore, at the subject-level, the precision mapping technique combined with ME-ICA enhanced the sensitivity in these regions, yielding bilateral activation in cortical, cerebellar, thalamic, and brainstem regions with both analyses. In addition, group-level and precision mapping regionspecific activation clusters were compared in standard and anatomical space. Results showcase the challenges of registration to a standard template brain, especially when dealing with variably located subcortical auditory nuclei across individuals, and the benefits of a wholebrain precision mapping approach which closely aligns an individual’s functional response to their anatomy. Lastly, precision mapping data were analyzed using SE and ME-ICA models and results were compared. Across subjects and regions, ME-ICA generally presented higher Dice similarity coefficients than SE starting at 20 minutes of data, particularly in thalamic and brainstem regions, suggesting that ME can provide great benefits in identifying small auditory nuclei in challenging regions with shorter scan times. Our results demonstrate compelling benefits of ME fMRI for studying the auditory system, particularly in combination with precision mapping of subcortical activation in individual subjects.

## Acknowledgements

This work was supported by the American Heart Association 25PRE1356822, NIH grants NIDCD K01DC019421, T32EB025766 and R03HD113915, the Center for Translational Imaging at Northwestern University, and through the computational resources and staff contributions provided for the Quest high performance computing facility at Northwestern University, which is jointly supported by the Office of the Provost, the Office for Research, and Northwestern University Information Technology. The authors would also like to thank Rachael Young for her support with data collection.

## Author Contributions

Michelle C. Medina: Conceptualization, Methodology, Software, Formal analysis, Investigation, Data curation, Writing - original draft, Writing - review & editing, Visualization, Project administration. Neha A. Reddy: Conceptualization, Methodology, Software, Formal analysis, Investigation, Writing - review & editing. Kevin R. Sitek: Methodology, Supervision, Writing - review & editing, Funding acquisition. Molly G. Bright: Conceptualization, Methodology, Supervision, Writing - review & editing, Project administration, Funding acquisition.

