## Supplemental Figure 1 for "Mapping Whole-Brain Auditory Activation with 3T Multi-Echo fMRI at the Group and Individual-Subject Level"

**Supplemental Materials**

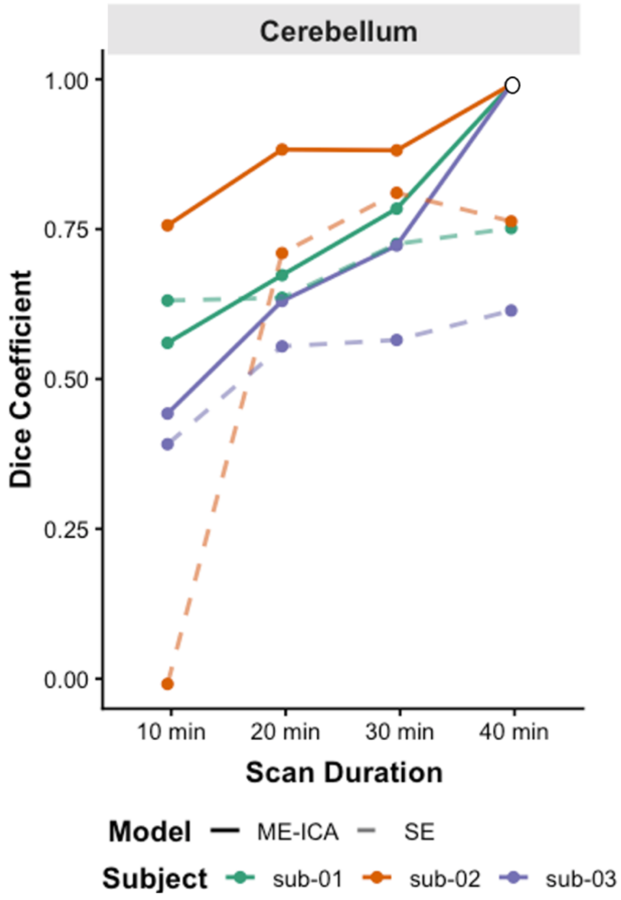


**Supplemental Figure 1:** Dice similarity coefficient plots between ME-ICA (solid line) and SE (dashed line) cerebellum activation results across scan duration. The 40-minute concatenated ME-ICA top 5% t-statistics cluster was used as ground-truth; thus, a blank circle was overlayed with each subject's 40-minute scan duration dice coefficient value. Each colored line represents a precision mapping subject.
